# Combining pangenomics and population genetics finds chromosomal re-arrangements, accessory-like chromosome segments, copy number variations and transposon polymorphisms in wheat and rye powdery mildew

**DOI:** 10.1101/2025.05.08.652814

**Authors:** Alexandros G. Sotiropoulos, Marion C. Müller, Lukas Kunz, Johannes P. Graf, Levente Kiss, Ralph Hückelhoven, Beat Keller, Thomas Wicker

**Affiliations:** Department of Plant and Microbial Biology, University of Zurich, Switzerland; Centre for Crop Health, University of Southern Queensland, Toowoomba, Australia; Chair of Phytopathology, TUM School of Life Sciences, Technical University of Munich, Freising, Germany

## Abstract

Grass powdery mildews (*Blumeria* spp.) include economically important fungal crop pathogens with complex and highly repetitive genomes. To investigate the diversity and genome evolution in *Blumeria graminis*, we combined population genetic and pangenomic analyses using a worldwide sample of 399 wheat powdery mildew isolates. Additionally, we produced high-quality genome assemblies for seven isolates from wheat and one from rye powdery mildew. Using these, we compiled the first grass powdery mildew pangenome comprising 11 *Blumeria graminis* isolates. We found multiple chromosomal rearrangements between the isolates that grow on wheat, rye and/or triticale hosts. Interestingly, chr-11 showed some characteristics of accessory chromosomes such as presence/absence of large chromosomal segments and higher sequence diversity. Additionally, we identified nearly 67,000 cases of copy number variations (CNVs), which were highly enriched within effector gene families. Furthermore, we found evidence for recent and high transposable element (TE) activity, such as high numbers of TE insertion polymorphisms. Analyses of TE families showed enrichment 1 kb to 2 kb up- and downstream of effector genes, and we also found high levels of TE insertion polymorphisms between populations. Our results demonstrate that chromosomal variations, gene family expansions and contractions, and TE activity are important sources of genome diversification and diversity in grass powdery mildews. Our findings indicate that a combination of pangenomic and population genetics analyses is needed to understand drivers of evolution in plant pathogenic fungi in a comprehensive way.

**Author Summary:** Fungi are a diverse group of organisms, with some of them (e.g. powdery mildews) having at times a devastating impact on important crops like wheat. Studying the genomes of these grass powdery mildew fungi can improve our understanding the pathogen’s patterns of genome evolution and virulence in order to find more efficient ways to protect crops. Using high quality genomes of a worldwide dataset of wheat and rye powdery mildews, we found multiple re-arrangements in chromosomes, as well as presence and absence of large chromosomal segments in different mildew strains. Along with some genes, important for making these pathogens infectious on various crop lines, some genomic regions may be deleted or duplicated, potentially affecting how the fungus survives and spreads. Additionally, we found that transposable elements are highly active in powdery mildews and that they are enriched near so-called effector genes, which are associated with fungal virulence. Our study shows how powdery mildew genomes have diversified during recent evolution, which has implications for future breeding of crops toward better resistance to fungal diseases.

## Introduction

Powdery mildews infecting cultivated and wild grasses include economically important plant pathogenic fungi, and those that infect crops can cause significant yield losses in agriculture (1). These pathogens are obligate biotrophs and obtain nutrients from the infected host plant tissues via the use of various effector proteins that modify the host’s metabolism. Earlier, all powdery mildews infecting diverse grasses (the Poaceae) were considered as belonging to a single species, named as *Blumeria graminis* (*B. graminis*) (2). Within this taxon, a number of taxonomically informal groups, known as formae speciales (ff. spp.), were delimited to identify distinct fungi that are each specialized to one or more species or genera of the Poaceae. For example, *B. graminis* f. sp. *tritici* referred to powdery mildews infecting wheat (*Triticum* spp.); *B. graminis* f. sp. *hordei* was used to identify powdery mildew on barley (*Hordeum vulgare*); and so on (3). A comprehensive host range testing revealed that some *Blumeria* isolates, identified as f. sp. *tritici*, infected tetraploid (durum) and hexaploid (bread) wheat varieties only; others grow better on tetraploid wheat varieties (4), and were included in the newly introduced f. sp. *dicocci*. Another newly identified group was named f. sp. *triticale* and includes isolates that infect triticale, an artificial hybrid crop between wheat and rye, as well as tetraploid and hexaploid wheat, and to a limited extent also some rye varieties (5). A recent taxonomic study recognised some of the formae speciales as different species of *Blumeria*; and retained the binomial *B. graminis* sensu stricto (s. str.) for powdery mildews characterised with distinct morphology and DNA barcode sequences that infect *Triticum*, *Secale*, and other species of the Poaceae (6). Our work follows the new taxonomy of *Blumeria* sensu Liu et al. (6), and refers to ff. spp. using the designations sensu (5,7), e.g., “*B.g. tritici*”, whenever needed for clarity.

During the past years, fungal genomics studies have become more feasible due to cheaper sequencing technologies. Studies on plant pathogenic fungi have highlighted differences in accessory chromosomes (e.g. in *Zymoseptoria tritici* and *Fusarium oxysporum*) (8,9), chromosomal re-arrangements (e.g. in *Cryphonectria parasitica*) (10), and Copy Number Variations (CNV) / Single Nucleotide Polymorphism (SNP) changes (e.g. in *Puccinia graminis*) (11). Powdery mildew genomic analyses were advanced by a number of high-quality genomes of multiple wheat and barley powdery mildew isolates that were recently determined (7,12–15). These highlighted the diversity of effectors in *Blumeria* spp. and expanded our knowledge on grass powdery mildews in general. Genomics of powdery mildew species infecting dicots, including grapevine, cucurbits, pepper, and a number of tree species (16–21), is less advanced compared to *Blumeria* spp. due, in part, to contamination and other quality issues detected in some genomes (22,23).

Transposable elements (TEs) are genetic components that can copy themselves and/or move around in the genome. There are two main classes of TEs: Class I (retrotransposons) and Class II (DNA transposons) (24). Retrotransposons tend to increase their copy numbers more rapidly than DNA transposons due to their copy-and-paste mechanism of replications (24). Long interspersed nuclear elements (LINEs), short interspersed nuclear elements (SINEs) and long terminal repeat (LTR) retrotransposons are widespread in the intergenic regions of many fungi (24,25). TEs have been well studied especially in humans, *Drosophila* and wheat. However, little is known about the TEs in fungi; studies conducted so far have revealed relatively lower TE content in most other fungal genomes compared to *B. graminis* (26). Furthermore, many fungi contain a specific pathway, i.e. repeat-induced point mutations (RIP), which disrupts TEs by introducing cytosine to thymine transitions (27,28), resulting in regions with extremely low G/C content, especially in non-genic, TE rich areas. Interestingly, *B. graminis* does not contain genes of the RIP pathway, and thus, its TEs are not affected by RIP (29). Possibly as a consequence of this, wheat powdery mildew has a high TE content (85% of its genome), among the highest known percentage of TE content in a fungal genome found to date (30). These two genomic patterns make *B. graminis* a good fungal model for studying TE diversity and evolution.

Pangenomes can be defined as the entire repertoire of genes accessible to the clade/species studied. The core genome contains genes present in all genomes, while accessory dispensable genes may be isolate or population specific (31). Pangenomic sequencing can help us study genome diversity on a species level by including more information about gene content, duplications and deletions of large genomic regions and other structural rearrangements. Sometimes, the addition of genes that were not found with short reads can be informative or differences in expression can be seen more clearly with various high-quality genomes. Various pangenomes of the hosts of *Blumeria* have been studied over the last few years such as that of hexaploid wheat (32) and barley (33). Similarly, there were studies on pangenomes of fungal model species (34) and model fungal plant pathogens e.g. *Zymoseptoria tritici* (35). With the advancement and improvements of long-read genome sequencing and assembling algorithms (36), along with more studies on the genomics of more *B. graminis* isolates (14,15,37), it has become more timely and feasible to study the pangenome of some other fungi with larger and more complex genomes, such as *B. graminis*.

Here, we present the analysis of a newly assembled grass powdery mildew pangenome based on near chromosome-scale assemblies of 11 isolates and re-sequencing data from about 399 isolates derived from various powdery mildew populations from across the globe. The following questions were addressed: (i) What are the levels of worldwide population genomic diversity in *B. graminis* f.sp. *tritici*, (ii) What are the structural and chromosomal differences in the genomes of different isolates, (iii) What is the extent of variation in content of effector and non-effector genes between isolates, (iv) Where are TEs located relative to genes in the *B. graminis* genome and how did TEs evolve in different isolates?

## Results

### New resources for the worldwide study of wheat powdery mildew populations

A dataset was compiled using a total of 399 isolates of *B. graminis* ff. spp. *tritici* and *dicocci* that were previously sequenced with Illumina short-read technology forming the basis of our analysis here (S1 Table). The dataset included the recently published Illumina short-reads of powdery mildew isolates from various parts of the world e.g. Egypt, Russia, USA, and the Fertile Crescent (7,14,38). The short reads produced in all these studies were used here and mapped to the reference isolate Bgt_CHE_96224 (13).

The assembled dataset included sequences of isolates from all continents, except Antarctica (Fig 1A). The majority of samples originate from the Fertile Crescent, which is the region of origin of both the wheat powdery mildew and its host (7). The Principal Component Analysis (PCA) of the SNP data (see methods) clearly revealed distinct populations (Fig 1B). Most of the variance in the PCA can be explained by the first two principal components, loosely reflecting latitude and longitude, mostly for the Eurasian wheat powdery mildew isolates (S1 Fig). The powdery mildew populations in North and South America, along with the ones in East Asia seem to be separated. The three isolates from Australia group together with the USA powdery mildew isolates as described before (7). The powdery mildew populations sampled in Europe, Egypt, the Fertile Crescent, central Russia and Kazakhstan overlap. This is in agreement with the hypothesis that there are traces of admixture between these populations, but also a gradient of geographic with genetic correlation (7).

**Fig 1.**
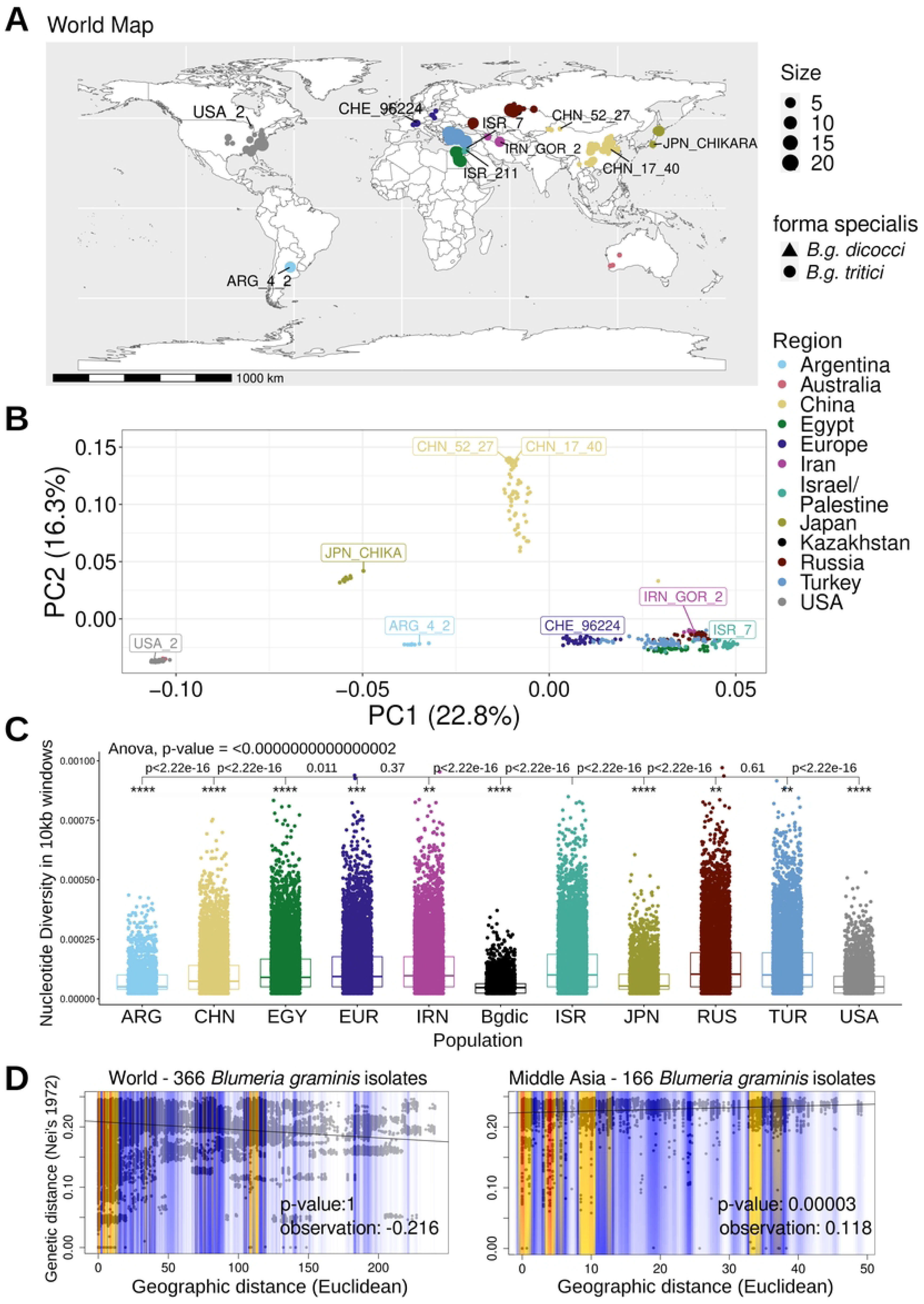
Worldwide dataset of wheat powdery mildew shows varying levels of genomic diversity and clustering populations. (A) A map of the isolates across the globe. (B) PCA of the SNPs of 399 isolates of *B. g. tritici* and *B.g. dicocci*. (C) Nucleotide diversity of the populations with at least eight isolates, based on eight randomly chosen isolates and analysing sequence windows of 10 kb, (D) Mantel test using the genetic data from the SNPs and the coordinates for the geographic correlation for a worldwide dataset where geographic information is available (on the left) and for the area around the region of origin (on the right).

Furthermore, nucleotide diversity of these populations (in 10 kb windows) (Fig 1C) shows that the populations that are more isolated in the PCA (i.e. ARG, USA, JPN, CHN) have a statistically less diverse mean. This can be explained by the isolated populations being further away from the region of origin with little genetic exchange. In this regard, they are similar to island populations, unlike the European – West Asian – North African populations, which hold the highest diversity even as individual populations. Furthermore, the *B.g. dicocci* population (Bgdic) has a low diversity which could be potentially explained by genetic bottlenecks resulting from small host populations as hypothesised before (7). A singletons analysis, identifying SNPs unique to only one isolate within a population, supported the results of the nucleotide diversity (S2 Table, S2 Fig). A mantel test indicated that there is no correlation between genetic distance and geography, while using all the global isolates. However, when using only the wheat powdery mildew isolates in the populations around the broader region of origin, which are not separated by any major geographic barriers, such as oceans or high mountains, there is a clear positive correlation (Fig 1D). This result supports a previous study that used a smaller dataset (7).

### A *Blumeria graminis* pangenome representing worldwide diversity

We sequenced eight *B. graminis* isolates from around the world previously collected (Fig 1A and 1B, S1 Table) with PacBio long-read technology, resulting in chromosome-scale assemblies. For this study, we also included chromosome-scale sequences of three previously published *B. graminis* isolates (CHE_96624, THUN-12, ISR_7) (13,38–40). We used a *B. hordei* genome (isolate DH14, GCA_900239735.1) (12) only when needed, due its fragmentation. While eight genomes belong to the forma specialis *tritici*, we also used one isolate of rye, triticale and tetraploid wheat mildews in the pangenomic dataset (S1 Table), resulting in a pangenome of a total of 11 powdery mildew genomes with reference genome quality similar to the reference Bgt_CHE_96224. The assemblies had genome sizes between ∼134Mb for the *B.g. secalis* isolate up to ∼143Mb for the *B.g. dicocci* isolate, with the *B.g. tritici* genome sizes falling in between these numbers. (S3 Table). The actual genome sizes differ, largely due to highly variable repetitive sequences in the centromeric region. As in a previous study (30), centromeres were defined by the presence of the centromere-specific retrotransposon family *RII_Fuji* (S3 Fig).

BUSCO (Benchmarking Universal Single-Copy Orthologs) analyses showed high levels of completeness of all chromosome-scale genome assemblies, with values above 94%, 97% and 98% completeness for the Leotiomycetes, the Ascomycota and the fungi reference dataset (“Fungi Odb10”), respectively (S4 and S5 Figs). The different genomes contain between 7,563 and 9,454 genes (S6A Fig), with the effector genes constituting ∼10% of the genes annotated, while the gene size distribution was consistent among isolates (S6B Fig). All newly sequenced isolates have 11 chromosomes as was found in the CHE_96224 reference isolate (S7A Fig and S3 Table) (30). Five isolates (CHE_96224, Bgtl_THUN-12, JPN_CHIKA, CHN_17_40, and CHN_52_27) include a few small contigs or a chromosome Unknown (chr-Un), where parts of the mitochondrial genome, along with un-anchored repetitive sequences, are collected. There is strong sequence collinearity between isolates, along all chromosomes, except in the highly repetitive centromeric regions (Fig 2). According to a phylogenetic network created with PhyloNet using 5,982 single-copy orthogroup proteins, *B.g. secalis* forms an outgroup together with the hybrid *B.g. triticale* isolate, which shows potential indication of recombination, also observed for the Bgt_JPN_CHIKA isolate (S7B Fig), as expected from previous studies (7).

**Fig 2.**
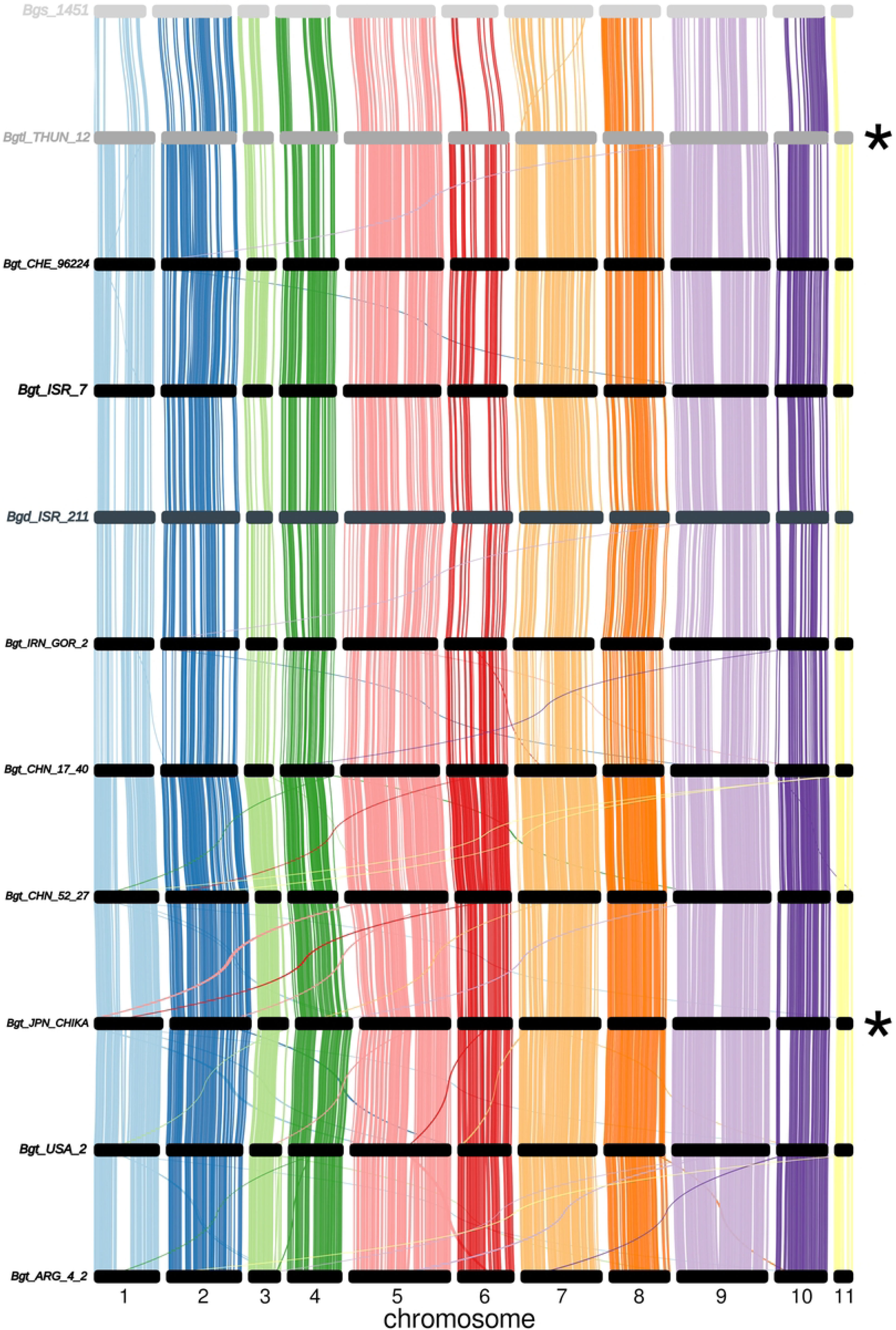
Chromosomal synteny of the 11 *B. graminis* isolates in the pangenome. The lines in colour indicate genomic regions of Minimum Alignment length of 100 kb. Each colour represents a chromosome from 1 to 11. The asterisks indicate the isolates from populations that were shown to be hybrids. The different shades of black and grey represent the different formae speciales. The fragmented *B. hordei* DH14 synteny with *B.g. tritici* CHE_96224 can be viewed in S8 Fig.

Average Nucleotide Identity (ANI) analysis in all chromosomes showed varying levels of homology: the *B.g. tritici* isolates share more than 99.45% homology with CHE_96224, while *B.g. dicocci* shares ∼99.34%, *B.g. secalis* ∼98.75%, and *B.g. triticale* ∼99.44% with the reference *B.g. tritici* CHE_96224 assembly within the isolates (S4 Table). This result reflects the evolution and phylogenetic distances of the respective isolates as expected.

The sequence of the mitochondrial genome for the isolate Bgt_CHN_17_40 was assembled in one contig (S9A Fig). The *B.g. tritici* mitochondrion genome was ∼104 kb in size and was also found completely in Bgt_chr-Un sequence (unordered sequence scaffolds that could not be assigned to nuclear chromosomes) in isolate CHE_96224. The mitochondrial genome sequence was also compared with the published *B. hordei* DH14 mitochondrial sequence (12). The *B.g. tritici* mitochondrial genome showed us high levels of collinearity with the published one from *B. hordei*, with ∼90 ANI sequence identity (S9B Fig and S4 Table) (12).

We also reviewed the mating type of these genomes, by using blast search of the published mating type loci on the genomes (S5 Table), using Bgt_CHE_96224 (MAT 1-2-1: *Bgt-3306* and SLA-2: *Bgt-2805*) (41), and the MAT 1-1-1 locus with the Bgt_CHN_52_27 gene *BgtCHN52_27_00382*. All the mating type loci were nearly identical between isolate and formae speciales with no more than one SNP difference. Both mating types were found in these 11 genomes, four isolates with MAT 1-1-1, and seven with MAT 1-2-1.

### Chromosomal and structural re-arrangements within *B.g. tritici*

An interchromosomal translocation was found between the isolate JPN_CHIKA (belonging to the Japanese wheat powdery mildew population) and the isolates USA_2 and CHN_52_27, which belong to the respective parental populations of the hybrid Japanese wheat powdery mildew population (Fig 3A) (7). This interchromosomal translocation affected a ∼640 kb segment containing 55 genes that was moved from chr-05 of CHN_52_27 to chr-01 of JPN_CHIKA, starting at ∼3.5Mb of chromosome 5. This segment corresponds to the left end of chr-01 in the JPN_CHIKA genome. This translocated region is also in chr-05 in the USA_2 genome (Fig 3A). Thus, we assume that it is a recent translocation in the JPN_CHIKA genome (S10 Fig). Furthermore, another translocation was found between chr-05 of CHN_52_27 and chr-02 of JPN_CHIKA, which amounts to approximately 242 kb and includes 27 genes (Fig 3A).

**Fig 3.**
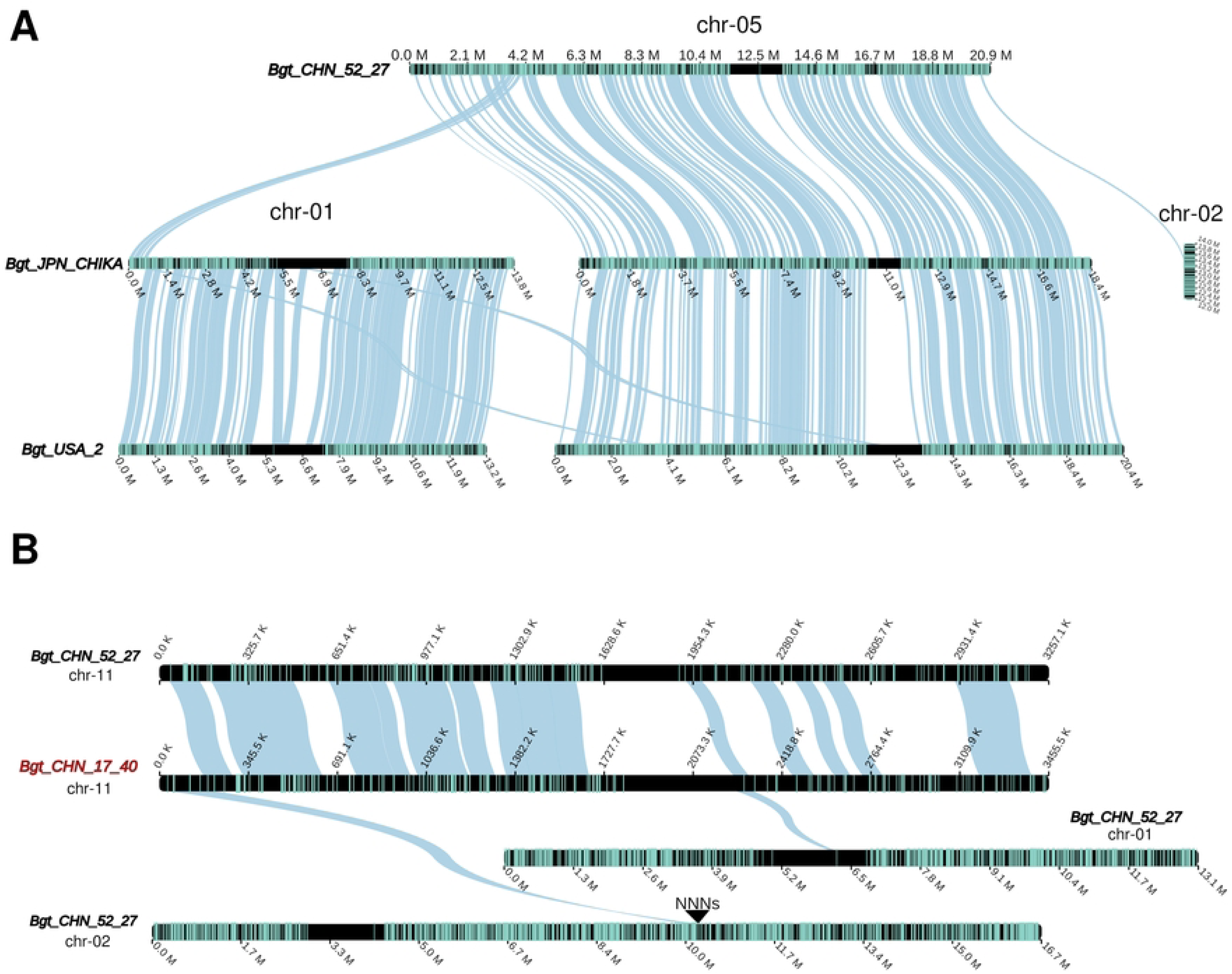
Examples of chromosome collinearity and re-arrangements within *B. g. tritici*. (A) Example of synteny and translocations in the genomes of three isolates (two coming from the parental populations, USA and CHN, and the middle genome coming from an isolate of the hybrid population of JPN powdery mildew. (B) Example of synteny in the genomes of two isolates from the same population (CHN powdery mildew). The petrol lines on the chromosomes indicate genes. The triangle and the “NNNs” show the position of a sequence gap.

It is well known that apparent re-arrangements (inversions, translocations) can be the result of mis-assemblies. We, therefore, examined the breakpoints of translocations for the presence of sequence gaps. We only found one gap next to a potential breakpoint (see below).

There are also re-arrangements between isolates coming from the same powdery mildew population (i.e. the Chinese powdery mildew isolates CHN_52_27 and CHN_17_40) (Fig 3B). In chr-11 of CHN_17_40, there are two segments with possible re-arrangements between chr-01 and chr-02 of isolate CHN_52_27. The one with chr-01 bears no sequence gaps (i.e. stretches of Ns) around the segment, while the one with chr-02 has a sequence gap on one side, which is why a possible miss-assembly cannot be excluded (in the position: 10,455,965 bp).

Such small structural re-arrangements between isolates are expected in a fungus with an annual sexual recombination cycle. However, more pronounced re-arrangements were found between different f. sp. isolates, as will be described hereafter.

### Chromosome 11 shows characteristics of an accessory chromosome

In *B.g. tritici* isolates, chromosomes 1 through 10 show generally high sequence conservation, while conservation is slightly lower between Bgt_CHE_96224 and Bgs_1459 (examples in S11 Fig and S4 Table), reflecting their phylogenetic distance (S7B Fig). However, striking differences were found in chr-11 between isolates of the different formae speciales (Fig 4). In general, sequence conservation of chr-11 is lower than in other chromosomes (examples in S11 Fig). Additionally, sequences on the right arm of chr-11 are less conserved than on the left arm (Fig 4A), both between *B.g. tritici* isolates and between *B.g. tritici* and *B.g. secalis*.

**Fig 4.**
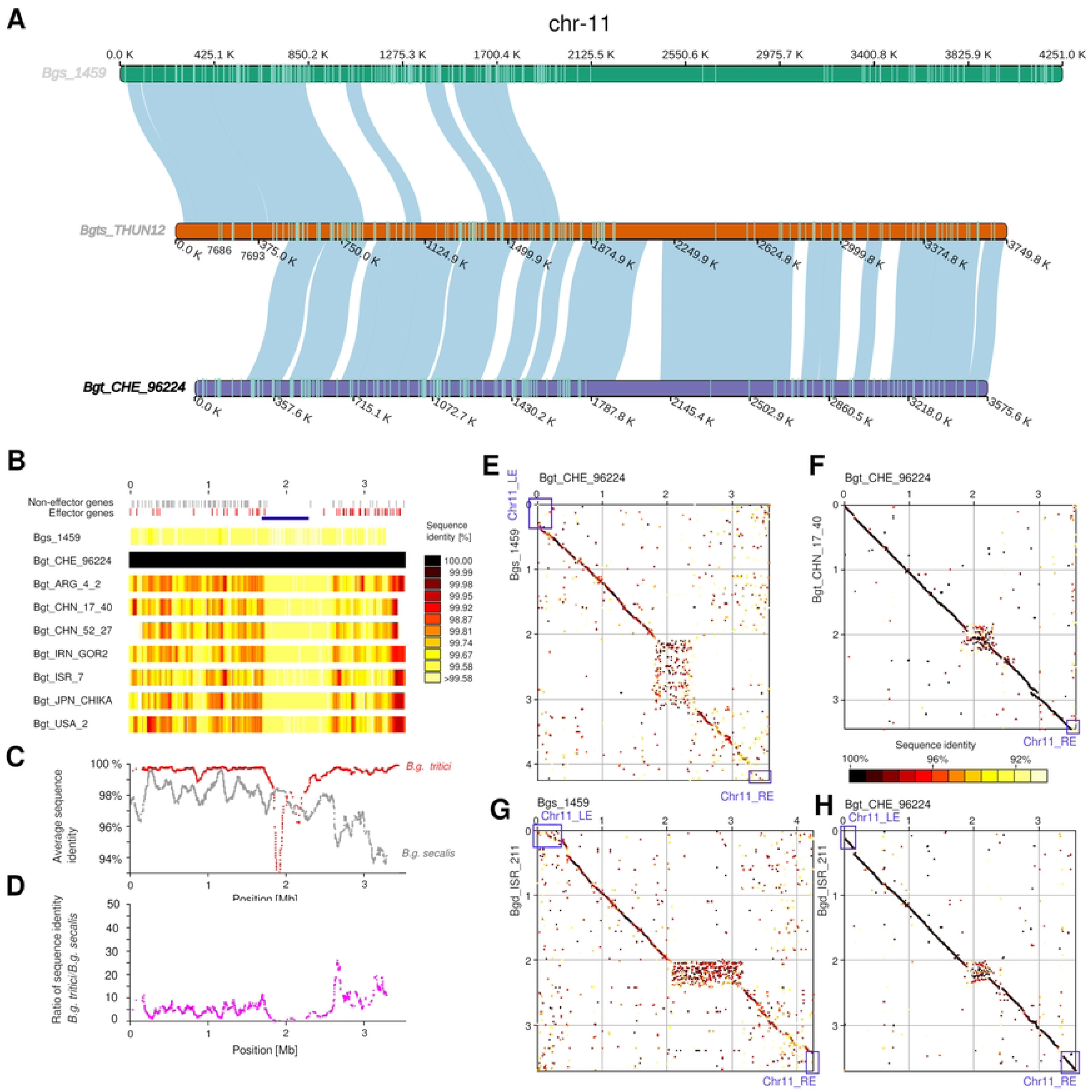
Analysis of structural variation and sequence diversity on *B. graminis* chr-11 indicates accessory chromosome-like characteristics. (A) Synteny in chr-11 for the hybrid *B. g. triticale* genome and its two-parent f. sp. genomes. The petrol lines on the chromosomes indicate genes. The colour of the names of the isolates corresponds to the forma specialis (black = *tritici*, darker grey = *triticale*, lighter grey = *secalis*). (B) Sequence conservation along chromosome 11. The heat map shows comparisons of *B. graminis* isolates with the reference isolate Bgt_CHE_96224 in 50 kb windows. Bgtl_THUN12 and Bgd_ISR_211 were excluded from this analysis. (C) The red line shows the average sequence conservation for all 50 kb windows among *B.g. tritici* isolates, while the gray line shows sequence conservation between Bgt_CHE_96224 and Bgs_1459. Note that sequence conservation is generally lower in the right chromosome arm. (D) Ratio of sequence conservation between *B.g. tritici* isolates and Bgs_1459. (E-H) Dot plot comparisons of chr-11 from *B. graminis* isolates. *B.g. secalis* and *B.g. dicocci* both have a segment at the left end (Chr11_LE, blue box) which is absent in *B.g. tritici*. Furthermore, a terminal segment of ∼100 kb is absent in Bgt_CHN_17_40, suggesting it contains accessory genes.

Noteworthy are the termini of chromosome 11: as previously described, the *B.g. triticale* genome is a recombined mosaic of the *B.g. tritici* genome (∼87,5%) and the *B.g. secalis* genome (∼12.5%) (5,39). In particular, in a previous study (39), the beginning of the left arm of chromosome 11 was found absent in *B.g. tritici* isolate Bgt_CHE_96224. Here, we studied this in more detail, and indeed found that the terminal ∼400 kb in Bgtl_THUN-12 originated from *B.g. secalis*, while the rest of the chromosome shows stronger synteny with *B.g. tritici*.

We found that the *B.g. secalis* and *B.g. dicocci* isolates both have this segment at the left end (Fig 4E, chr11_LE), while it is absent in all *B.g. tritici* isolates. Conversely, an ∼100 kb segment at the right end of chr-11 is absent in *B.g. secalis* when compared to *B.g. tritici* (Fig 4E, Chr11_RE). Interestingly, this segment is present in the *B.g. dicocci* isolate (Fig 4G-4H), suggesting *B.g. dicocci* might represent the ancestral state as it has both the segment on at the left and right end of chr-11. Furthermore, a terminal segment of ∼100 kb on the right end of the chromosome is absent in Bgt_CHN_17_40 (Fig 4F, Chr11_RE). This segment is also absent in Bgt_CHN_52_27 (S12A Fig). Additionally, isolate Bgt_CHN_52_27 is missing a ∼180 kb segment on the left side of chr-11 (S12A Fig), while Bgt_ARG_4_2 has an extra segment of about ∼150 kb on the left side compared to Bgt_CHE_96224 (S12B Fig). We, therefore, propose that the termini of chr-11 contain accessory genes, which may be involved in host specificity.

We produced a genealogic tree of all the predicted candidate effector proteins found on the right side of the chromosome arm of the Bgt_CHE_96224, Bgd_ISR_211 and Bgs_1459 isolates (S13 Fig). We also included the effectors found in the exclusive non-*B. g. tritici* left chromosome arm segment. Most of these genes belong to the previously described large effector family E003 (30). Furthermore, we included effector proteins from chr-06 that had homology with candidate effector *Bgs1459-09640*, a gene that is found on the left end segment in *B.g. secalis*. This group of effector genes has apparently seen an expansion specifically in *B.g. dicocci* (S13 Fig). Furthermore, a gene *Bgs_09664* from the left end of chr-11 of *B.g. secalis*, groups with homologs from the right end of chr-11 in *B.g. tritici* (S13 Fig), indicating that the ancestral chr-11 had E003-family effectors on both ends, which underwent differential loss in the different formae speciales (with 28 E003-family effectors in chr-11 for Bgt_CHE_96224, 20 for Bgs_1459 and 33 for Bgd_ISR_211).

Finally, three additional smaller segments which originate from *B.g. secalis* in *B.g. triticale* were found in the ∼1.6 Mb at the left end of chr-11 (Fig 4A). This segment contains a gene that is only found in *B.g. secalis* and *B.g. triticale*, but is absent in *B.g. tritici* (S1 Note, S14 Fig).

We hypothesize that presence/absence of chromosome segments, differential gene losses and family expansions of candidate effectors in chr-11 may be among the factors that contribute to host specificity.

### A variety of gene duplications and deletions in the *B. graminis* pangenome

The structural variations in chr-11 inspired us to further study presence and absence of genes in the pangenome. Here, we used short read sequence coverage data to identify possible gene copy number variants (CNVs). If coverage with sequence reads was particularly high for a gene in a given isolate (i.e. roughly multiples of the average sequence coverage), we took this as evidence for multiple copies of that gene. Using a normalized coverage cutoff of 0.1 < coverage < 1.7, we identified 66,856 individual CNVs for the various isolates (Fig 5A, S15 Fig). We observed that the number of duplicated genes was statistically significantly higher in a few populations (e.g. EGY population) and between formae speciales (Fig 5A). For example, *B.g. secalis*, has approximately 2.5 times more gene duplications than the *B.g. tritici* isolates, while *B.g. dicocci* has approximately 1.5 times more. These results could be attributed to the larger genomic differences between the different formae speciales, but also to using CHE_96224 as the reference genome, which creates a reference bias and possible underestimate of CNV frequencies. Furthermore, even within *B.g. tritici* isolates, we found that populations from the greater Fertile Crescent area were among the most diverse (Fig 5A).

**Fig 5.**
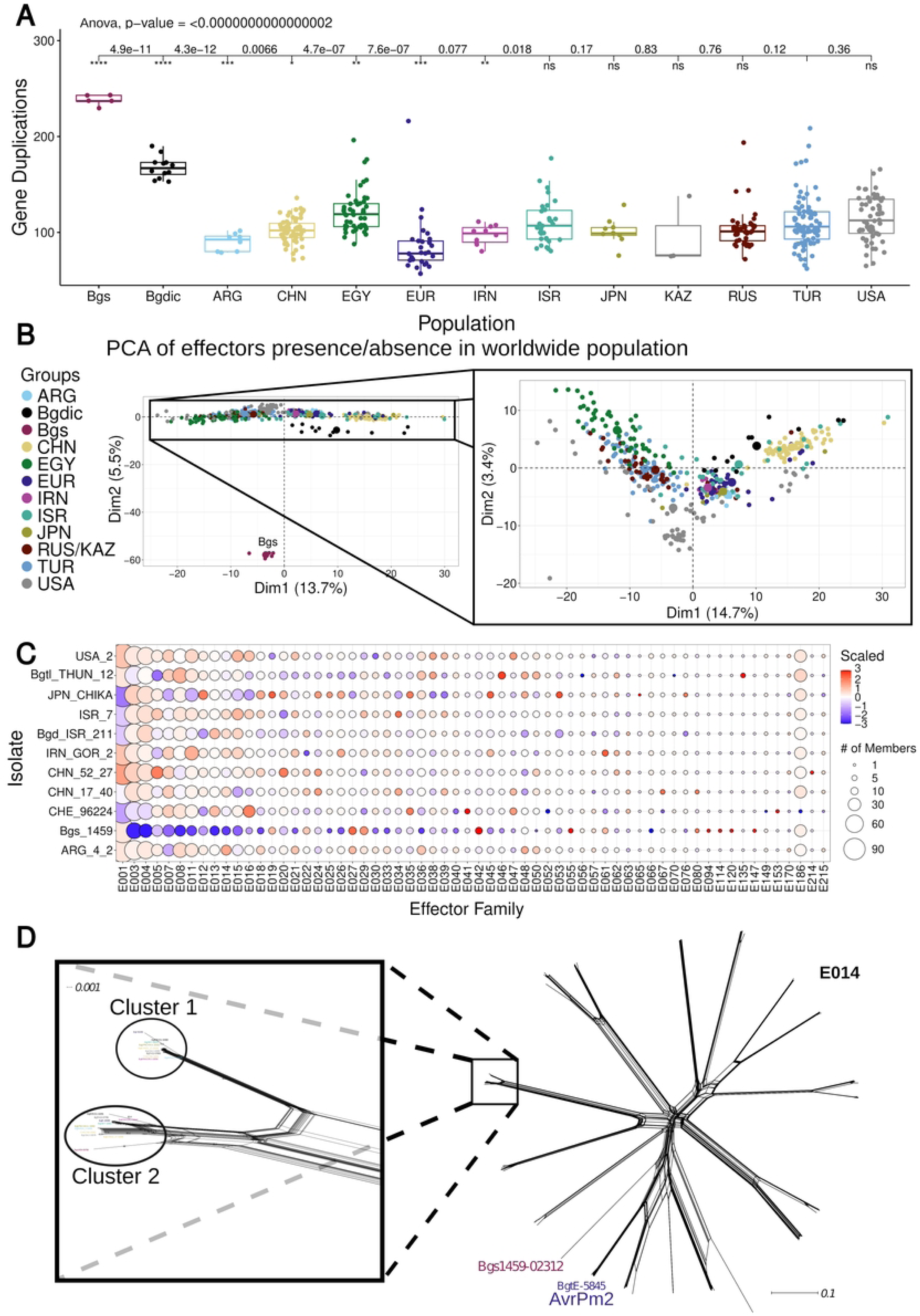
Duplications/Deletions of genes using the populations and the pangenome. (A) Number of gene duplications per population. The asterisks signify statistical significance of difference between other populations and the TUR powdery mildew population. (B) PCA of effector gene presence/absence summary per isolate, including the *B.g. secalis* isolates (left) and without them (right). (C) Expansion or shrinkage of effector families in the pangenome. Number of effector genes per family in all 11 *Blumeria graminis* isolates. (D) Phylogenetic network of one of the most diverse effector families in wheat powdery mildew (E014).

We identified 3,795 genes with indications for CNV in at least one isolate. We created a PCA from a matrix of sequence coverage of each effector gene per isolate for all isolates including the *B.g. secalis* ones (Fig 5B, left). The latter isolates clearly differ from the rest of the *B. graminis* isolates. If we exclude the *B.g. secalis* isolates, the PCA (Fig 5B) shows that even though most populations cluster separately (e.g. *B.g. dicocci*), they seem to overlap. Additionally, further examination revealed differences between populations and gene families (S16 Fig), especially in the case of effector genes (S16A Fig). Furthermore, *B. graminis* isolates other than *B.g. tritici* tend to have higher number of CNVs as expected for more distant isolates.

We then searched the individual genomes to identify potential gene enrichments in specific isolates (Fig 5C). The *B.g. secalis* isolate 1459, even though it seems to have gene deletions in many of the effector families, also shows expansions of a few effector gene families (e.g. E027, E029, E042, E055, E080, E094, E114, E120, E147, Fig 5C). Furthermore, THUN-12 has an exclusive expansion of effector gene family E135, where THUN-12 has four effectors, while the other 10 isolates have only one (Fig 5C). Additionally, the isolate JPN_CHIKA has a specific expansion of family E012 (JPN_CHIKA has 17 members, while others have between 11 and 13) (Fig 5C). Also, for some isolates of the same powdery mildew populations (CHN), there can be expansions in one isolate for one family whilst not in the other (i.e. E001 and E005).

An interesting effector family is E014, with the identified avirulence gene *AvrPm2* as a member (42). Family E014 has more members in some isolates (e.g. ARG_4_2), while isolate Bgs_1459, has a reduction of gene number. Despite its smaller number of genes, Bgs_1459, has some effectors in this family that are unique and do not have homologues in the other isolates (i.e. Bgs1459-02312), as shown in a phylogenetic network of this family for all the pangenome isolates (Fig 5D).

### Transposable Elements insertions correlate with host specialisation and divergence of individual powdery mildew populations

Since TEs have been shown to play an important role in powdery mildew evolution (30) and in host-pathogen interactions (43,44), we analysed the TE composition in the *B. graminis* pangenome, as well as in seven of the wheat powdery mildew populations (ARG, CHN, *B.g. dicocci*, EUR, IRN, ISR and USA populations). All 11 isolates that are part of the pangenome have similar repeat content of approximately ∼55% of the genome (S6 Table) and similar abundance of TE superfamilies (S17 Fig). DNA (class 2) transposons have around 8.5% of annotated copies from the ∼21,000 annotated and classified TEs on average per isolate (Fig 6A). SINEs (Short Interspersed Nuclear Elements) account for about 38% of the annotated TEs. LINEs (Long Interspersed Nuclear Elements) account for about 14%, while different LTR retrotransposon superfamilies account for more than 40% of annotated TEs.

**Fig 6.**
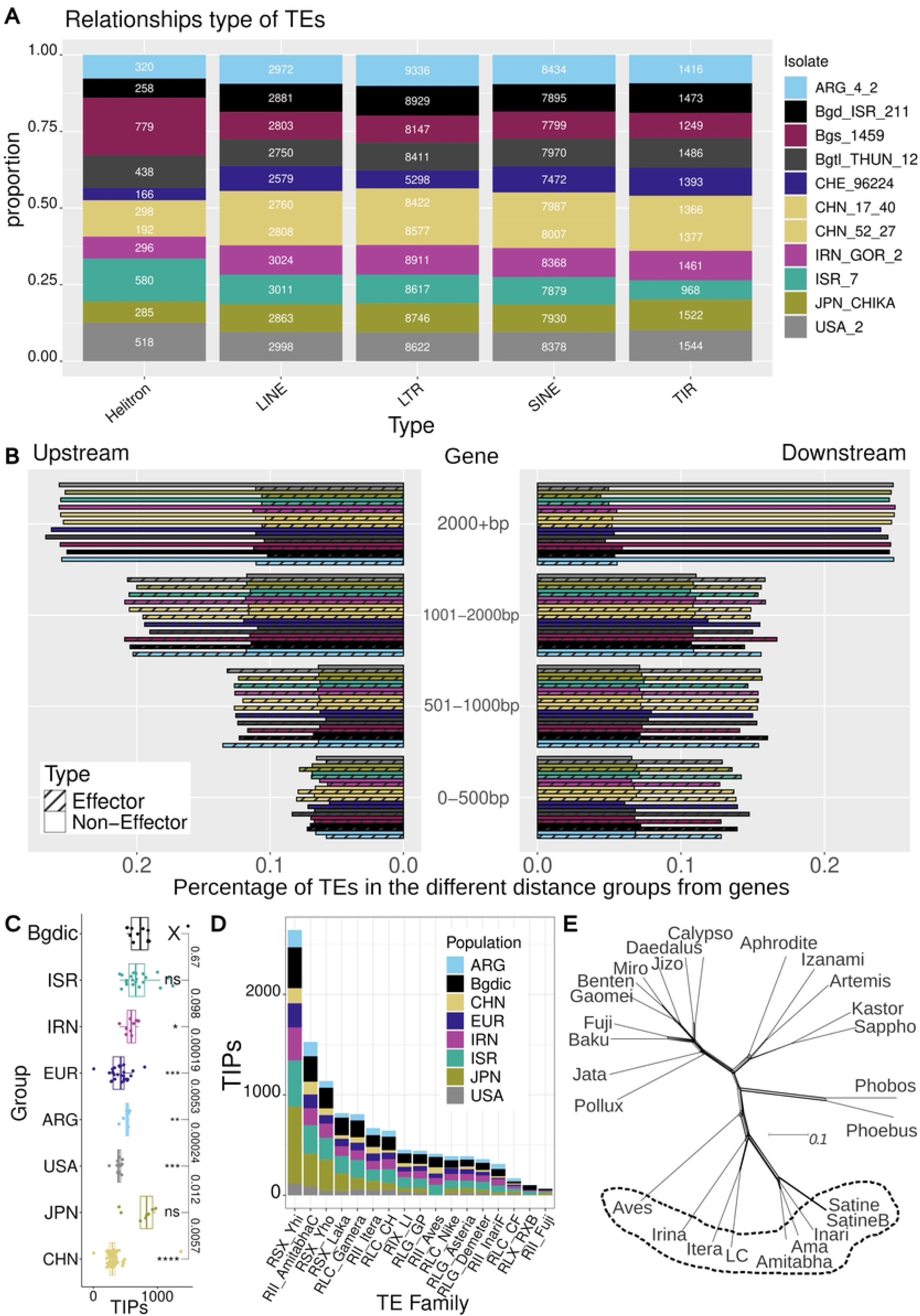
TE analyses in wheat powdery mildew. (A) Annotated copies of TE types in the different isolates of the pangenome. (B) Number of all TEs found in various distance ranges downstream and upstream of genes for all the isolates of the pangenome, see legend of Fig 6A. The bars with the dotted lines refer to distance from effector genes, while the ones without any lines refer to the non-effector genes. (C) Boxplot of the number of TIPs per isolate for populations with eight or more isolates (isolates with less than 2000 TIPs are shown, extreme outliers were excluded, because they were interpreted as technical artifacts). The “X” symbol refers to the population that the statistical comparisons have been made, were the “ns”, “*”, “**”, “***”, and “****” show significant difference with said population. (D) Analyses of TE insertions (TIPs) in the different wheat powdery mildew populations normalized for sample size for each population. Depicted are the 17 TE families that showed most insertion polymorphisms in all populations. (E) Phylogenetic network of the alignment of reverse transcriptase (RT) proteins for most of the LINE TE elements, using consensus sequences from isolate CHE_96224. The TEs in the circle include the three LINEs with the most TIPs in the populations tested.

We found an enrichment of Helitrons in the *B.g. secalis* isolate, which has 779 annotated copies, compared to the other isolates which have between 166 and 580 (Fig 6A). Moreover, the hybrid *B.g. triticale* isolate that originates from Europe has an intermediate number of Helitrons compared to its ancestors *B.g. secalis* Bgs_1459 and *B.g. tritici* CHE_96224. We then analysed TE composition at different distances up- and downstream of genes (Fig 6B). The majority of TEs can be found at a distance greater than 1,000 bp away from a gene, with less than 3,000 TEs (out of the approximately 21,000 on average per isolate) found closer than 500 bp to a gene (Fig 6B). Interestingly, TEs are closer to the effector genes, with particular enrichment of TEs around 1-2 kb distance from the effector genes.

We used the software detettore (45) to detect transposon insertion polymorphisms (TIPs) using the paired end information of Illumina read mappings of isolates to a reference genome. We ran detettore on mappings of 166 *B. graminis* isolates against the isolate Bgt_CHE_96224 (see methods and S7 Table). The populations closer to the region of origin, along with the *B.g. dicocci* population had the most TIPs (Fig 6C). The CHN powdery mildew population had the least TIPs which can be attributed to the shorter Illumina reads, posing a known bias. Furthermore, the hybrid JPN population has the most TIPs, something that could be hypothesized that it has to do with its hybridisation.

When looking at specific TEs, SINEs and LINEs were found to be the most mobile TEs in the various wheat mildew populations with the *RSX_Bgt_Yhi* and *RII_Bgt_Amitabha_C* families being the two with the most polymorphic insertions observed across populations (Fig 6D). The three populations in the Fertile Crescent, along with the hybrid powdery mildew population from Japan, showed the highest numbers of TIPs for most families. After LINEs and SINEs, LTR retrotransposons are the ones with most polymorphic insertions, particularly the families *RLC_Bgt_Gamera* and *RLC_Bgt_CH*. The top three LINE families with most TIPs against Bgt_CHE_96224, were found across the genome (S18 Fig). An interesting TE family, *RLX_Bgt_RXB* had higher numbers of TIPs in the *B.g. dicocci* population compared to all the other populations studied.

Many factors such as evolutionary pressure or genetic drift could have led to this increase. While it is tempting to speculate that the high number of lineage-specific TE insertions may also have driven the host-specificity of *B.g. dicocci*, more studies are needed to support this hypothesis.

Although both SINEs and LINEs generally showed high TIP numbers, only LINEs encode a conserved protein (reverse transcriptase). This allowed the study of their relative divergence time. For this, we used the predicted reverse transcriptase proteins from multiple LINE families of the Bgt_CHE_96224 isolate. The proteins were used for a multiple sequence alignment with ClustalW and construction of a phylogenetic network with PhyloNet and splitstree (Fig 6E). Some LINE proteins clustered together and were also the families with the most TE insertions (e.g. *RII_Bgt_Amitabha_C*, *RII_Bgt_Itera* and *RII_Bgt_Aves*). This indicates that the LINE cluster that includes these TEs (Fig 6E, in dotted circle) could have evolved more rapidly compared to other LINEs and could also include some of the most active LINE TEs in these genomes.

## Discussion

Various pangenomic studies on fungal plant pathogens have highlighted the genomic diversity of various species, like accessory chromosomes in the hemibiotrophic fungus *Zymoseptoria tritici* that can increase virulence in different isolates (35), or accessory chromosomes in the hemibiotroph *Fusarium oxysporum* that indicate host specificity (46). Little is known on pangenomic level for most obligate plant biotrophic fungi like in powdery mildews. Studying the synteny of near-chromosome scale assemblies together with population genetics of short-read sequences for an economically important and well-studied obligate plant pathogen like *Blumeria graminis*, has helped us better understand its genome diversification.

Recently, *B. graminis* was re-defined from a taxonomic point of view, and seven additional species were delimited within the genus *Blumeria* based on their host ranges, DNA barcode sequences, and morphology (6). Our work has highlighted the importance of using formae speciales sensu Menardo et al. (5) and Sotiropoulos et al. (7) to clearly distinguish grass powdery mildews with distinct host ranges and genomic patterns within the recently defined *Blumeria graminis* species.

There are many facets of fungal genome architecture that play a role in the genome evolution. One of them is the presence of accessory chromosomes which could play a role in the evolution of gene clusters in filamentous some fungi such as in *Fusarium* spp. and *Alternaria* spp.(47). *B. graminis* has not been known to have any accessory chromosomes in a strict sense (i.e. presence/absence of entire chromosomes), which has been also supported from the research here. Another facet of structural genome diversity are chromosomal re-arrangements. Large structural re-arrangements could contribute to reproductive isolation, and therefore, to host specialisation. Even though we observed multiple re-arrangements in this study, we did not notice any big structural rearrangements (e.g. chromosome arm relocation), especially within *B.g. tritici* isolates. However, chr-11, the smallest chromosome in *B. graminis*, has the highest number of variation and some of the largest re-arrangements. This is reminiscent of characteristics of accessory chromosomes. Indeed, our study shows particular variation at the ends of chr-11 with large chromosomal segments being present or absent between formae speciales and even within *B.g. tritici*.

Furthermore, we found numerous CNVs of genes in the *B. graminis* populations. We identified a difference in CNV frequency in genes between effectors and non-effectors on average, with a higher CNV ratio in effectors (∼0.58) compared to non-effectors (∼0.38), despite no specific differences in CNV frequency within individual populations, apart from the isolates that did not belong to *B.g. tritici*.

Moreover, the transposable elements landscape can have an impact on genome evolution, especially in species where most of the genome comprises of TEs like this one (13,41). We found many TE polymorphisms in the *B. graminis* pangenome, especially between formae speciales. Studying TE polymorphisms showed an overall high and recent activity of TEs in *B. graminis*, which goes in line with the divergence time of these species. Previous studies indicated that *B. graminis* diverged at about 10 Ma ago from *B. hordei*, a couple less Ma compared to each host *Triticum* spp. diverging from *Hordeum* spp. (48). Within the *B. graminis* species, *B.g. tritici* seems to have diverged from other formae speciales (such as *B.g. secalis*) much more recently (less than 300,000 years) than its grass hosts (with rye and wheat diverging at about 4 Ma) (49). We also found that TEs are generally closer upstream of effector genes than in non-effector genes, which is puzzling. One possible explanation would be that effector genes simply have shorter promoters and thus allow close-by TE insertions. Another explanation would be that TEs favor open chromatin for insertion, with promoter regions of effector genes probably representing the regions with the most accessible DNA within the genome, as effectors consistently are among the highest expressed genes (CITE 4 papers). Alternatively, relaxed selection in effector genes could allow for this to happen. Finally, TEs near effectors could potentially provide templates for unequal homologous recombination which could facilitate copy number variation and/or emergence of recombinant genes.

In general, we showed that pangenome data is suited for identifying structural changes and large-scale chromosomal re-arrangements. Furthermore, the inclusion of short read sequencing data helped identify CNVs and presence/absence of genes and TEs in populations, even in the absence of chromosome-scale assembly for a particular isolate. These types of polymorphisms are often overlooked since most studies only focus on SNPs. Thus, we conclude that our hybrid approach of using chromosome-scale assemblies and re-sequencing data is well suited to identify different types of polymorphisms. Adding sequencing data from more isolates in the future can provide an even clearer view of the genomic diversity and evolution of *B. graminis*.

## Materials and methods

### Fungal isolates collection

The whole-genome sequenced isolates were introduced into the collection by reviving the ascospores in chasmothecia (50) following the methods of (30) (S1 Table). The isolates that were acquired through chasmothecia were single-cell colony isolated to ensure that only one clonal individual was collected. Each isolate was propagated on ten-day-old freshly cut wheat leaves (cultivar “Kanzler”) in Petri dishes. After four days, we picked with a toothpick a single colony to pass to new fresh leaves. We repeated this twice to make sure we have isolated one individual and then proceeded to propagate it, in order to get many spores for sequencing. The dates of collection of the isolates vary between 1990 and 2019 and the isolates are listed in the S1 Table. For each region, isolates have been collected from various fields and various cultivars from *Triticum aestivum* and *T. turgidum*. A list of the isolates, the region, the coordinates of collection, the year of collection and the wheat species/subspecies that they were collected from is shown in the S1 Table. As outgroups for some analyses the forma specialis *B. g. secalis* was used.

### DNA preparation and whole-genome sequencing

The DNA extraction method that was used, was done with a chloroform- and CTAB based-protocol modified from Bourras and colleagues (51). Illumina paired-end reads sequencing of 150 bp read length and an insert size of ca. 500 bp was performed to generate 1 to 6 Gb sequence data per isolate using NovaSeq 6000 technologies. The rest of the isolates were sequenced before with the same extraction protocol and their accessions were found on SRA. The Illumina sequence data is accessible from the NCBI Short Read Archive (the project numbers are shown in S1 Table).

### Read mapping and variant calling

The raw Illumina reads were trimmed for adapter contamination and better sequencing quality. For trimming, the software Trimmomatic V0.38 (52) was used with the following settings: illuminaclip=TRuSeq3-PE.fa:2:30:10, leading=10, trailing=10, slidingwindow=5:10, minlen=50. The reference genome of *B.g. tritici* CHE_96224 version 3.16 was used to align the reads by using the short read aligner bwa mem v0.7.17-r1188(53) with the following settings: -M. We used samtools v1.9 (54) for sorting and removing duplicate reads. In order to mark PCR duplicates in the alignment (bam) files, we used the MarkDuplicates module of Picard tools version 2.18.25-SNAPSHOT. The average genome-wide coverage (calculated as the number of mapped reads multiplied by average mapped read length and divided by the reference genome size) ranged between x10 to x60 for all the isolates. We finally used ‘samtools index’ to create indices for the bam files. The mapped reads were also further checked visually for their quality and GC% content, using the software qualimap2 and the multi-bamqc function (55). Isolates that were found to be clonal in a previous study where excluded (7) leading to 399 isolates used for most analyses.

The mapped reads were used to make variant calling files using GATK v.4.1.2.0 (56–58). We started by using the HaplotypeCaller with the options: --java-options “-Xmx4g”, -ERC GVCF, -ploidy 1. Afterwards, we used CombineGVCFs to combine the single gvcf files into one file and then, with GenotypeGVCFs we performed joint variant calls on the merged variant gvcf file using the option: -- max-alternate-alleles 4. We used the GATK VariantFiltration function to hard filter SNPs with quality thresholds recommended by GATK (57,58) and used with other plant pathogenic fungi (59,60). The following thresholds were used: QUAL < 450, QD < 20.0, MQ < 30.0, −2 > BaseQRankSum > 2, −2 > MQRankSum > 2, −2 > ReadPosRankSum > 2, FS > 0.1. Then, we filtered for a genotyping rate (>99.9%) and removed the indels. Finally, we kept only biallelic SNPs using vcftools: --max-alleles 2 (version 0.1.5) (61). The number of SNPs retained was 60,258.

### Population genomics analyses

To select the isolates for sequencing we performed various population genetics analyses. We used R (v4.3.3) (62), and the R packages tidyverse (v2.0.0) (63), ggplot2 (v3.5.1) (64), and stats (v4.3.3) (62) for many of the following analyses. Inkscape (v1.3.2) was used to compile many figures (65). We first created a map with the samples using R packages: sf (v1.0-18) (66,67), rnaturalearth (v1.0.1) (68), ggspatial (v1.1.9) (69), and ggrepel (v0.9.3) (70). Then, we made a PCA using vcftools (v0.1.16), and visualised the results in R. Furthermore, we used packages and ggplot2, ggpubr (v0.6.0) (71), and gdistance (v1.6.4) (72) to look at the diversity of the populations. We used again vcftools to identify the singleton SNPs (SNPs found only in one isolate/individual within a whole population) once using all isolates and another time with only eight random isolate per populations. Finally, we used packages vcfR (v1.14.0) (73), StAMPP (v1.6.3) (74), gplots (v v3.2.0) (75), MASS (v v7.3-60) (76), grDevices (v4.3.3) and graphics (v4.3.3) (62) to study the genetic and geographic correlation between the isolates. The package rcartocolor (v2.1.1) was used to find colour-blindness friendly colours used in many of the plots (77).

### PacBio sequencing, gene and TE annotation

In addition to the Illumina sequencing, we long-read sequenced with the PacBio technology eight powdery mildew isolates (six *B.g. tritici*, one *B.g. dicocci* and one *B.g. secalis*). There was 1x SMRT Cell 1M reads used per isolate, apart from isolate CHN_17_40, where 8M reads with PacBio HiFi were used. The reads from the isolates were then initially assembled using SMRT Link software tools (for versions see in S8 Table) at the Functional Genomics Center Zurich (FGCZ) in Switzerland. The assembled contigs were then scaffolded into near chromosomes by aligning them to the reference genome of isolate CHE_96224, Bgt_genome_v3_16 (13). Each contig was placed and orientated by its best blastn hit (first: e-value, second: bit-score). Contigs with no blast hit were gathered into a chromosome Unknown. Contigs were merged with stretches of 200 “N” insertions (sequence gaps). We also used the already sequenced and assembled genomes of two *B.g. tritici* (CHE_96224, ISR_7) and one *B.g. triticale* (THUN-12) isolate for the pangenome analyses (S9 Table). One recently sequenced genome in the European wheat powdery mildew population (isolate CHVD_042201) was not used (78), as it was not accessible at the time this analysis was done. The assembled genome data is accessible from NCBI (the project numbers are shown in S1 Table). The difference between these genome was evaluated using the OrthoANIu tool (v1.2), a standalone average nucleotide identity (ANI) calculator (79).

The gene annotations were performed using maker (v2.31.10). The annotations were generated by using the Bgt_CDS_v4_23 proteins as a template. Maker was used with the prot2genome function and the repeats and transposons were found with maker internal TE_proteins, as well as the initial annotations in PTREP and nrTREP databases (https://trep-db.uzh.ch/). On a second maker round, the annotation was created based on the *B. hordei* gene annotations of isolates DH14 and RACE1 (GCA_900239735.1 & GCA_900237765.1). From the resulting genes in the second round, only genes with no annotation in their corresponding loci in the first round (non-redundant genes) were added to the final annotation (as extra genes). In order to assess the genome and assemblies and annotations we used BUSCO (v5.4.4) (80,81) along with some further analyses below. We looked at the size of chromosomes and created a phylogenetic network using Splitstree4 (version 4.19.1, 27 Jun 2023) (82), and 100 number of replicates for bootstrapping, with the orthogroups within *B. graminis* (S7 Fig) via Orthofinder (v2.5.5) and the parameter “-msa” (83). The number of genes and their type (effector, non-effector etc) was visualised, along with the protein size distribution to further check for the quality of the annotation (S6 Fig).

TEs were further annotated by first using the TE database for *Blumeria graminis* nrTREP (version 24, https://trep-db.uzh.ch/) and the software EDTA (v.2.0.1) (84). We then manually curated newly identified TEs and performed a final annotation with the latest *Blumeria* repeats deposited in nrTREP (S10 Table). The mitochondrial genome sequences were identified by using blastn of the mitochondrial contig of the *B. hordei* DH14 mitochondrial genome to the genome assembly of isolates CHE_96224 and CHN_17_40. For the former, pieces of the mitochondrial genome were scattered in Bgt_chr-Un which we eventually did not use, while for the latter, the mitochondrial genome was in one contig. We then, used the software MITOS2 (v2.1.9) to annotate the one mitochondrial genome (85). The annotated output was visualised using OrganellarGenomeDRAW (v1.3.3) (86).

### Synteny in the pangenome

In order to study any structural re-arrangements in the *B. graminis* pangenome, we looked at the synteny between the chromosomes and the various isolates with a Minimum Alignment Length set to 50 kb (for the individual chromosomal re-arrangement visualisation in Fig 3A-3B) or 100 kb (for the overall chromosomal synteny in Fig 2, and Fig 4A). For the synteny between the isolates *B.g. tritici* CHE_96224 and *B. hordei* DH14 (S8 Fig), the Minimum Alignment Length was set to 10 kb, since these genomes are more dissimilar than the genomes within the *B. graminis* species. We used software NGenomeSyn (v1.41) to perform this analysis (87). We used the Paul Wong colour accessible palette the various isolates and populations across the rest of the study (88).

### Analysis of sequence conservation between chromosomes

To study sequence conservation between isolates along chromosomes, 1 kb segments of the reference isolate Bgt_CHE_96224 were used in blastn searches against the corresponding chromosomes of all other isolates. Blastn hits were then filtered for collinearity, requesting that a given 1 kb segment had its top blastn hit within 5% of its position in the reference chromosome. With this, we filtered out non-specific matches stemming from repetitive sequences. Sequence conservation between Bgt_CHE_96224 and a given isolate was then calculated as a running average over 50 of the colinear 1 kb segments. Overall sequence conservation across all *B.g. tritici* isolates was then calculated as the average of all 50 kb windows in the *B.g. tritici* isolates. This was used to produce the conservation plots shown in Fig 4C and S11 Fig. Finally, the ratio of sequence conservation between *B.g. tritici* isolates, and *B.g. secalis* was calculated for all windows, resulting in the plots in Fig 4D and S11 Fig. Chromosome-scale tot plots were produced by running blastn searches of 1 kb segments of a chromosome of one isolate against the corresponding chromosome of another. Positions of the hits in both chromosomes were recorded and used for the dot plots shown in Fig 4.

Finally, to study phylogeny of candidate effectors from chr-11 and selected homologs from other chromosomes, we created a dataset with all the candidate effector identified proteins from the right chromosomal arm of Bgs_1459, Bgd_ISR_211, and Bgt_CHE_96224, along with the effectors on the left end region that is missing in the *B.g. tritici* isolates and homologous effectors of Bgs1459-09640 found in chr-06 after using blast (v2.7.1+) (89) on these three genomes to find any homologous protein. We excluded three proteins that had a size smaller than 50 amino acids (three were excluded). We added the unclassified candidate effector protein Bgs1459-00747 as an outgroup. We aligned the dataset with ClustalW (v.2.1) (90,91), and constructed a phylogenetic tree using the Bayesian inference with MrBayes (v.3.2.7a) (92,93) with settings nst=6 and rates=invgamma. The Markov chain Monte Carlo (mcmc) analysis was run until the probability value decreased to under 0.02 with a sampling frequency of 10 and a burn-in of 25% of samples. We visualised the tree with Figtree (v1.4.4) (94) and we used colours from Paul Tol’s accessible palettes (https://sronpersonalpages.nl/~pault/).

### Identification of specific candidate effectors

The study of some specific effectors of interest was made by first changing the annotation files gff to gtf format using gffread (v0.12.8) (95). We then used STAR (v2.5.2a) (96) to map already published RNAseq data (namely SRR6410427, a replicate of RNAseq data of isolate Bgtl_THUN12 on the Timbo triticale line 2 days post infection). Using IGV (v2.15.4, 12/08/2022) (97), we manually checked the expression and the annotation of specific genes BgTH12- and Bgs1459-0. After using blast (v2.7.1+) (89) to find the closest homologous proteins on the whole pangenome and on the *B. hordei* isolate DH14, we aligned them with ClustalW (v.2.1) (90,91) and visualised the alignment with Jalview (v2.11.1.4+dfsg-3) (98). A phylogenetic tree was constructed using the same method as described above.

### Identification of gene duplications and deletions

We used the Illumina DNA mapped reads and the annotation v4_20 to calculate read coverage of genes using the logic of the software featureCounts (v2.0.0) (99) in order to identify duplications and deletions of genes. We normalized the Illumina read coverage result of each gene to the average coverage of all genes per respective chromosome. The normalized coverage of each gene per isolate was divided by the coverage of the reference isolate CHE_96224. Subsequently, genes displaying normalized coverage below 0.1 were considered deleted, and genes showing coverage above 1.7 were considered duplicated.

### Analyses of transposable elements for the pangenome

We analysed positions of TEs relative to genes with the TEGRiP pipeline (github: https://github.com/marieBvr/TEs_genes_relationship_pipeline) (100,101). In order to find TE insertions which are polymorphic between isolates, we used the software detettore (v.2.0.3) (45,102) and a subset of isolates (S7 Table). We used ClustalW (v.2.1) (90,91) to produce the alignments of the TE consensus sequences of families of interest, and then changed the format to nexus and checked the alignment with ClustalX (v2.1) (91,103). We used MrBayes (v.3.2.7a) with settings nst=6 and rates=invgamma to infer trees (92,93). The Markov chain Monte Carlo (mcmc) analysis was run until the probability value decreased to under 0.01 with a sampling frequency of 10 and a burn-in of 25% of samples. We visualised the tree using FigTree (v1.4.4) (94).

## Acknowledgments

The authors would like to acknowledge Dr. Coraline Praz and Dr. Manuel Amos Poretti for the discussions with the authors regarding powdery mildew transcriptomics. This project was mainly supported by grants from the University of Zurich Research Priority Program. This project was also partly supported by the Centre for Crop Health of the University of Southern Queensland, through the Discovery Project DP210103869 funded by the Australian Research Council.

